# Estrogen represses *Tgfbr1* and *Bmpr1a* expression via estrogen receptor beta in MC3T3-E1 cells

**DOI:** 10.1101/170084

**Authors:** Hanliang He, Chunqing Wang, Qifeng Tang, Fan Yang, Youjia Xu

## Abstract

MC3T3-E1 is a clonal pre-osteoblastic cell line derived from newborn mouse calvaria, which is commonly used in osteoblast studies. To investigate the effects of estrogen on osteoblasts, we treated MC3T3-E1 cells with various concentrations of estrogen and assessed their proliferation. Next, we performed RNA deep sequencing to investigate the effects on estrogen target genes. *Bmpr1a* and *Tgfbr1*, important participants in the TGF-beta signaling pathway, were down-regulated in our deep sequencing results. Bioinformatics analysis revealed that estrogen receptor response elements (EREs) were present in the *Bmpr1a* and *Tgfbr1* promoters. Culturing the cells with the estrogen receptor (ER) alpha or beta antagonists 1,3-bis(4-hydroxyphenyl)-4-methyl-5-[4-(2-piperidinylethoxy)phenol]-1H-pyrazole dihydrochloride (MPP) or 4-[2-phenyl-5,7-bis(trifluoromethyl) pyrazolo[1,5-alpha]pyrimidin-3-yl] phenol (PTHPP), respectively, demonstrated that ER beta is involved in the estrogen-mediated repression of *Tgfbr1* and *Bmpr1a*.The chromatin immunoprecipitation (ChIP) results were consistent with the conclusion that E2 increased the binding of ER beta at the EREs located in the *Tgfbr1* and *Bmpr1a* promoters. Our research provides new insight into the role of estrogen in bone metabolisms.

## Introduction

Estrogen, a steroid hormone in the human body, plays an important role in many physiological processes [1, 2], such as the maintenance of secondary sexual characteristics [3], the development of neurons [4], and bone homeostasis [5]. Estrogen is critical to maintain bone and bone density [6]. Estrogen deficiency often leads to the emergence of osteoporosis. In postmenopausal women, estrogen secretion is markedly reduced, as is bone mineral density, whereas the incidence of bone fractures is increased. This form of osteoporosis is called postmenopausal osteoporosis (PO) [7]. An imbalance of osteoblasts and osteoclasts in the bone causes osteoporosis [8]. PO has become one of the most important diseases that affects older women, as it entails huge financial burdens for individuals and society, and reduces the quality of life [9].

In 1988, Komm et al. found that ERs are expressed in osteoblasts [10], suggesting that estrogen may have a regulatory role for osteoblasts. After the discovery of ERs on osteogenic precursors and bone cells, an increasing number of studies has shown that estrogen has a direct effect on osteogenesis, as well as an indirect effect that does not depend on ERs. Kousteni et al. and others have found that estrogen can inhibit osteoblast apoptosis and prolong osteoblast life, thereby increasing the osteogenic capacity of each osteoblast [11]. The effect of estrogen is achieved by rapid activation of ER-mediated regulatory kinases [12]. The lack of estrogen causes a marked increase in NF-κB activity in osteoblasts [13]. Estrogen deficiency leads to an imbalance in bone metabolism between bone formation and resorption, eventually leading to osteoporosis. As the mechanism is not yet clear, further studies on estrogen regulation of bone metabolism and its signaling pathways are required to achieve a better understanding.

MC3T3-E1 cells are a clonal pre-osteoblastic cell line derived from newborn mouse calvarias, which is utilized in many osteoblast studies [14]. To investigate the effects of estrogen on osteoblasts, we treated MC3T3-E1 cells with various concentrations of estrogen (17-β estradiol, E2), then monitored their proliferation and apoptosis; we then conducted RNA deep sequencing to investigate estrogen target genes. From the differentially expressed genes (DEGs) in the estrogen-treated groups compared with the DMSO control, we chose *Tgfbr1* and *Bmpr1a* as candidate genes, as they play important roles in the transforming growth factor beta (TGF-β) and bone morphogenetic protein (BMP) signaling pathways. Promoter analysis revealed estrogen receptor response elements (ERE) in the *Tgfbr1* and *Bmpr1a* promoter regions. Using the ERα or ERβ antagonists 1,3-bis(4-hydroxyphenyl)-4-methyl-5-[4-(2-piperidinylethoxy)phenol]-1H-pyrazole dihydrochloride (MPP) or 4-[2-phenyl-5,7-bis(trifluoromethyl) pyrazolo[1,5-α]pyrimidin-3-yl] phenol (PTHPP), respectively, demonstrated that ERβ is involved in the repression of estrogen in the regulation of *Tgfbr1* and *Bmpr1a*.The chromatin immunoprecipitation(ChIP) results were consistent with the conclusion that E2 increased binding to the EREs located in the *Tgfbr1* and *Bmpr1a* promoters. Thus, we conclude that estrogen affects bone through negative regulation of *Tgfbr1* and *Bmpr1a* in MC3T3-E1 cells. Our research provides new view insight into the mechanisms of action of estrogen on osteoblasts.

## Materials & Methods

### Cell culture

The clonal murine pre-osteoblastic cell line MC3T3-E1 was used in this study. The cells were thawed from frozen stocks and cultured in 100-mm culture dishes in α-MEM supplemented with 10% (v/v) fetal bovine serum (Gibco), 1% (w/v) antibiotics/antimycotics (the stock solution contained 100 U/mL penicillin G sodium, 100 μg/mL streptomycin sulfate, and 0.25 μg/mL amphotericin B, in saline), 1.0 mM sodium pyruvate, 0.1 mM nonessential amino acids, and 1.5 g/L sodium bicarbonate (Gibco) at 37°C in a humidified atmosphere containing 5% (v/v) CO_2_. After establishing cultures from frozen cells, we sub-cultured the cells several times. Confluent cells were detached using 0.25% trypsin in Mg^2+^- and Ca^2+^-free phosphate-buffered saline (PBS) before use. For the treatment of MC3T3-E1 with E2, the cells were seeded in each well of a 24- or 6-well plate with 0.5 or 2mL of culture medium. The cells were incubated in the presence of E2 at different concentrations (0.01, 0.1, 1, and 10nM), and each concentration of E2 was maintained for three days. A group that received DMSO served as the control.

### Cell apoptosis assay

Cell apoptosis of MC3T3-E1 cells was analyzed with the TUNEL Apoptosis Detection Kit (Beyotime). Briefly, the cells were collected two days after treatment with E2, washed with PBS, and suspended in 500μl binding buffer. Then, the cells were stained for the detection of apoptosis, according to the manufacturer’s instructions. The ratio of apoptotic cells was analyzed under an optical microscope by counting terminal deoxynucleotidyl transferase dUTP nick end labeling (TUNEL) positive cells in six randomly selected areas.

### Total mRNA extraction

Total cell mRNA was extracted using TRIzol^®^, according to manufacturer instruction (Invitrogen). Smart Spec Plus (Bio-Rad) was used to measure the absorption at 260/280 nm to assess the quality and quantity of the collected RNA. Lastly, the integrity of the extracted RNA was further assessed using 1.5% agarose gel electrophoresis. Subsequently, the RNA was transcribed to first strand cDNA using the First Strand cDNA Synthesis Kit (Takara) for gene expression analysis.

### RNA sequencing ibrary construction and sequencing

We used 20μg of total RNA from the control group and the10 nM E2-treated group for the RNA sequencing (RNA-seq) library preparation. The collected mRNA was purified and concentrated using oligo (dT)-conjugated magnetic beads (Invitrogen) before library preparation. The purified mRNA was randomly cut into fragments with the fragmentation buffer. We used the mRNA as a template, with 6-base random primers (random hexamers), to synthesize the first strand of cDNA. We then added dNTPs, RNase H, and DNA polymerase I to the buffer, to synthesize the second strand of cDNA. Lastly, we used AMPure^®^ XP beads (Beckman) to purify the synthesized cDNA. The purification of double-stranded cDNA was performed with end repair and A-tailing. The AMPure^®^ XP beads were used to select the size of the fragments. Finally, the cDNA library was obtained by PCR. Using “sequencing by synthesis” technology, we sequenced the cDNA libraries with the Illumina HiSeq2500 high-throughput sequencing platform and obtained high-quality reads. The reads and bases sequenced by the platform were usually considered raw data, and most of them got a Q30 score for base quality.

### Analysis of DEGs

Gene expression has time-and space-specific characteristics, and both external stimulation and the internal environment can affect gene expression. Under two different conditions (such as the control and treatment, wild-type and mutant, different times, different organizations, etc.), genes that have significantly different expression are considered DEGs. Similarly, transcripts with significantly different expression are called differentially expressed transcripts. In bioinformatics, the process of finding differentially expressed transcripts or DEGs is known as differential expression analysis.

### Annotation and analysis of DEGs

The Gene Ontology (GO) database was built in 2000, and is a standard, structured biological annotation system aimed at establishing a system of standard vocabulary and knowledge of genes and their products. The GO annotation system contains three main branches: biological process, molecular function, and cellular component. We used the GO database to predict the functions of the DEGs identified in this study[15]. Enrichment analysis of the differences between samples was carried out using Top GO. Top GO directly displays the GO node and the hierarchical relationship of the DEGs. The Clusters of Orthologous Groups (COG) of proteins database is constructed on the basis of the phylogeny of bacteria, algae, and eukaryotes to classify the orthologues of gene products. To annotate the pathways of the DEGs, it is necessary to analyze the functions of the genes. The Kyoto Encyclopedia of Genes and Genomes (KEGG) database is a major metabolic pathways database[16]. The analysis of the over-presentation of the DEGs in a pathway is important to determine the functions of the genes. In our study, COG and KEGG were used to analyze the DEGs.

### Quantitative real-time PCR (qRT-PCR) validation

Total cell mRNA was extracted using TRIzol^®^, according to manufacturer instruction (Invitrogen). We subjected 10μg of total RNA to DNaseI treatment with 1 U DNaseI (NEB). The reaction was carried out at 37°C for 10 min followed by heat inactivation at 65°C for10 min. We then used 2.5μg of DNase-treated RNA for cDNA synthesis with reverse transcriptase (Bio-Rad), in accordance with the manufacturer’s protocol. The expression of the *Gapdh* gene, which the transcriptome database indicated was stable, was used as the control for qRT-PCR experiments. Primers were designed for selected transcripts from the transcriptome database and real-time PCR was performed with SYBR^®^Green I master mix (TAKARA) on a CFX Connect ™ Real-Time PCR Detection System (Bio-Rad). The relative expression of the transcripts was calculated using the Δ Δ Ct method. The primers are shown in Table 1.

**Table 1.**
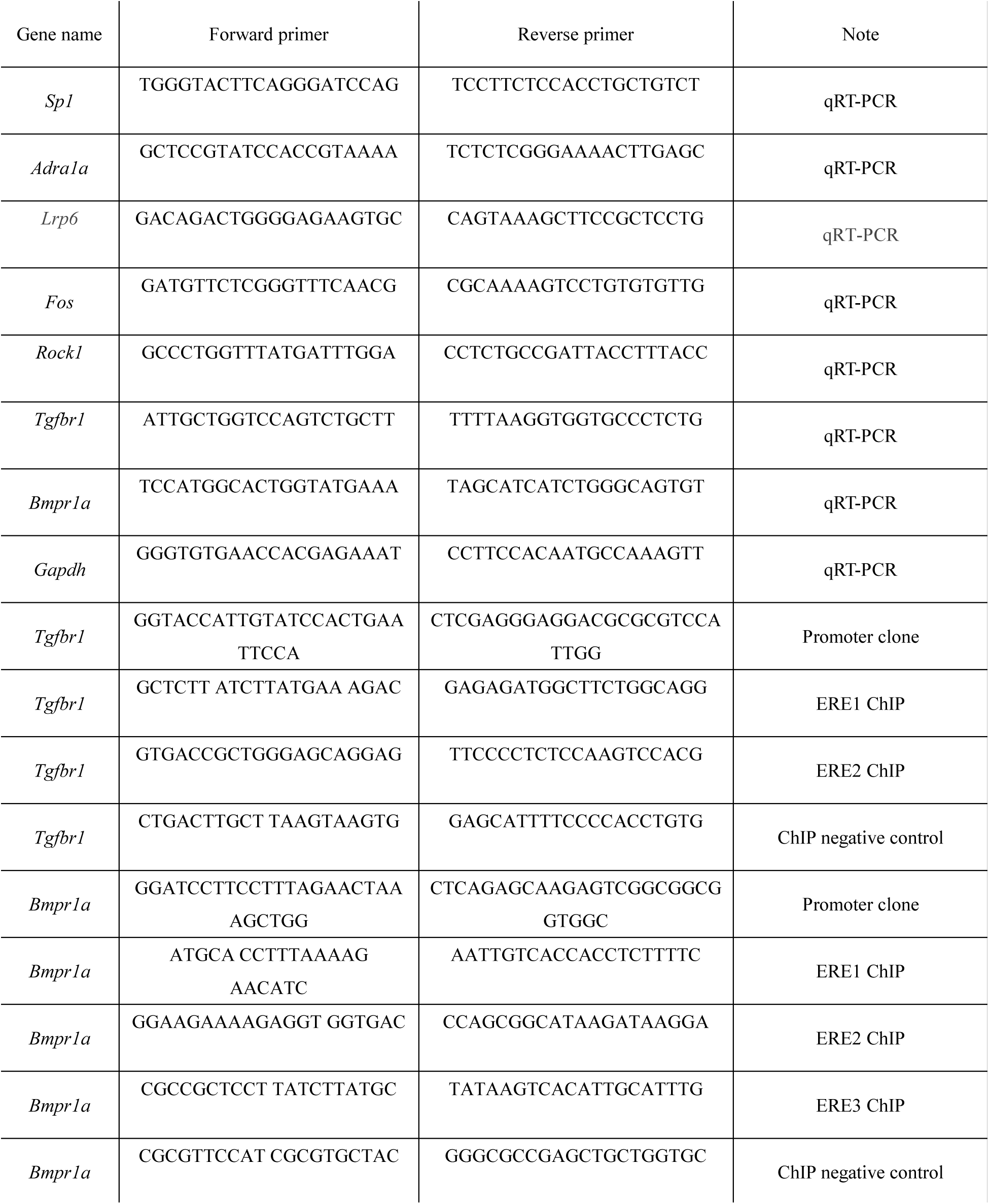
The primer of list in this paper

### ERα and ERβ antagonist treatment of MC3T3-E1 cells

For the ERα and ERβ antagonist experiments, the ERα antagonist MPP and the ERβ antagonist PTHPP were added to the MC3T3-E1 cultures. After 72 h of treatment with the antagonists, the cells were harvested to quantify the target gene expression.

### Chip assays

ChIP assays were performed according to the manufacturer’s protocol (ChIP Assay Kit, Millipore). Briefly, MC3T3-E1 cells were collected after culture with or without E2, cross-linked in 2% formaldehyde at 28^0^C for 30 min, then treated with a 1/10 volume of 1.25 M glycine to stop cross-linking, followed by PBS washes (three washes for 10 min each). We used purified rabbit or mouse IgG (Invitrogen) as a negative control and an antibody against ERβ to pull down the DNA. We performed ChIP PCR using primers flanking the ERE sites, as well as primers not flanking the ERE sites in the promoter regions of *Tgfr1* and *Bmpr1a* as controls. The primers used for the ChIP PCR are listed in Table 1.

### Statistics

Data are presented as mean± standard deviation. Statistical differences between two groups were determined with Student’s t test. Statistical differences among groups were analyzed with a one-way ANOVA followed by Student’s test. All experiments were repeated at least three times, and representative experiments are shown. Differences were considered significant at P<0.05.

## Results

### E2 increased MC3T3-E1 cell proliferation

To determine if estrogen affects MC3T3-E1proliferation, we treated MC3T3-E1 cells with different concentrations of E2 (0, 0.01, 0.1, 1, and 10nM E2). We then performed CCK-8 experiments. We found that MC3T3-E1 cell proliferation was unchanged in the low concentration E2-treated groups (0.01 and 0.1 nM E2) compared with that in the DMSO control groups (Fig. 1). However, in the high concentration E2 groups (1 and 10nM E2), MC3T3-E1 cell proliferation was significantly higher than in the DMSO control groups (Fig. 1). These data revealed that high concentrations of E2 promote MC3T3-E1 cell proliferation. In order to obtain robust data and explore the mechanisms of action of estrogen on MC3T3-E1 cells, we next analyzed cells treated with the highest, supraphysiological concentration of E2 (10 nM) by RNA-seq, to explore the mechanisms of action of estrogen on MC3T3-E1 cells.

**Figure 1.**
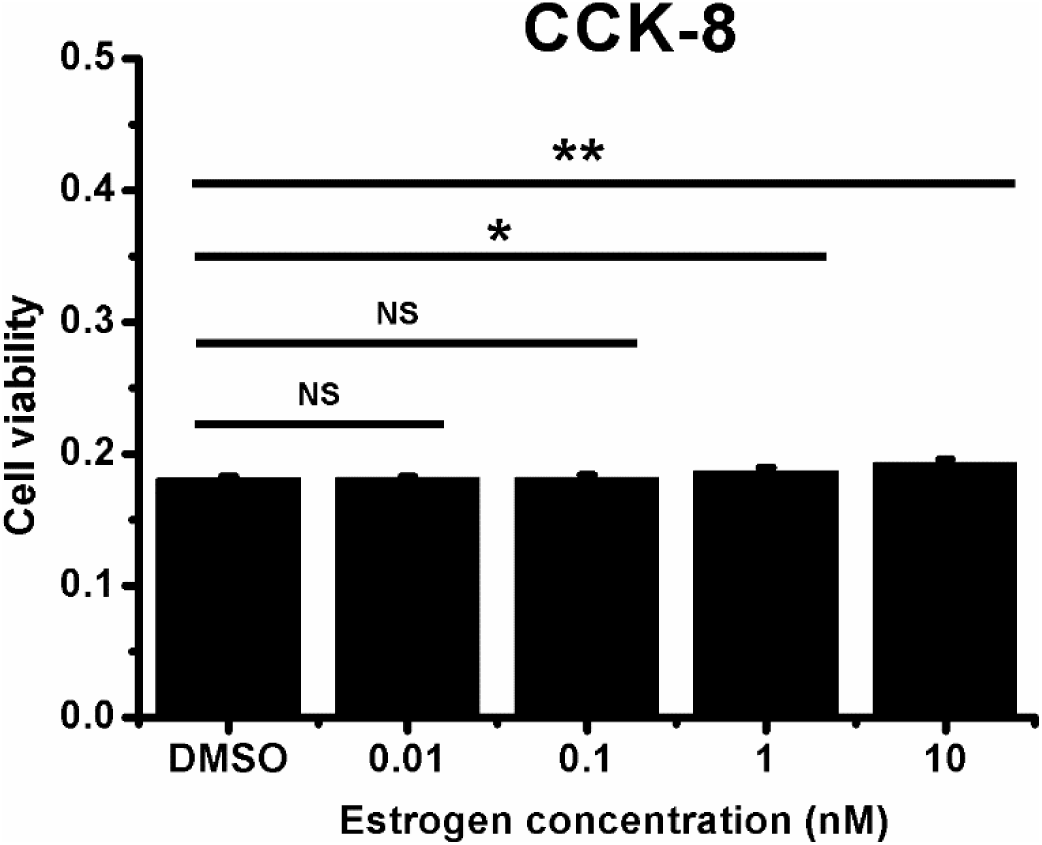
The effects of estrogen on MC3T3-E1 proliferation. MC3T3-E1cells were treated with different concentrations of E2, then subjected to analysis with a CCK-8 assay. Student’s t-test. ^∗^, P<0.05; ^∗∗^P<0.01.

### RNA-seq and identification of DEGs

To assess the effects of E2 on gene transcription, we used the Cuffdiff analysis module Cufflinks to analyze the differential gene expression in the samples [17]. The screening criteria for significant differences in the expression of genes are |log2Ratio| ≥ 1 and q-value≤ 0.05. Using these criteria, we identified 460 DEGs. Among those DEGs, 66 genes were up-regulated and 394 genes were down-regulated in the cells treated with 10 nM E2 treated group (Fig. 2).

**Figure 2.**
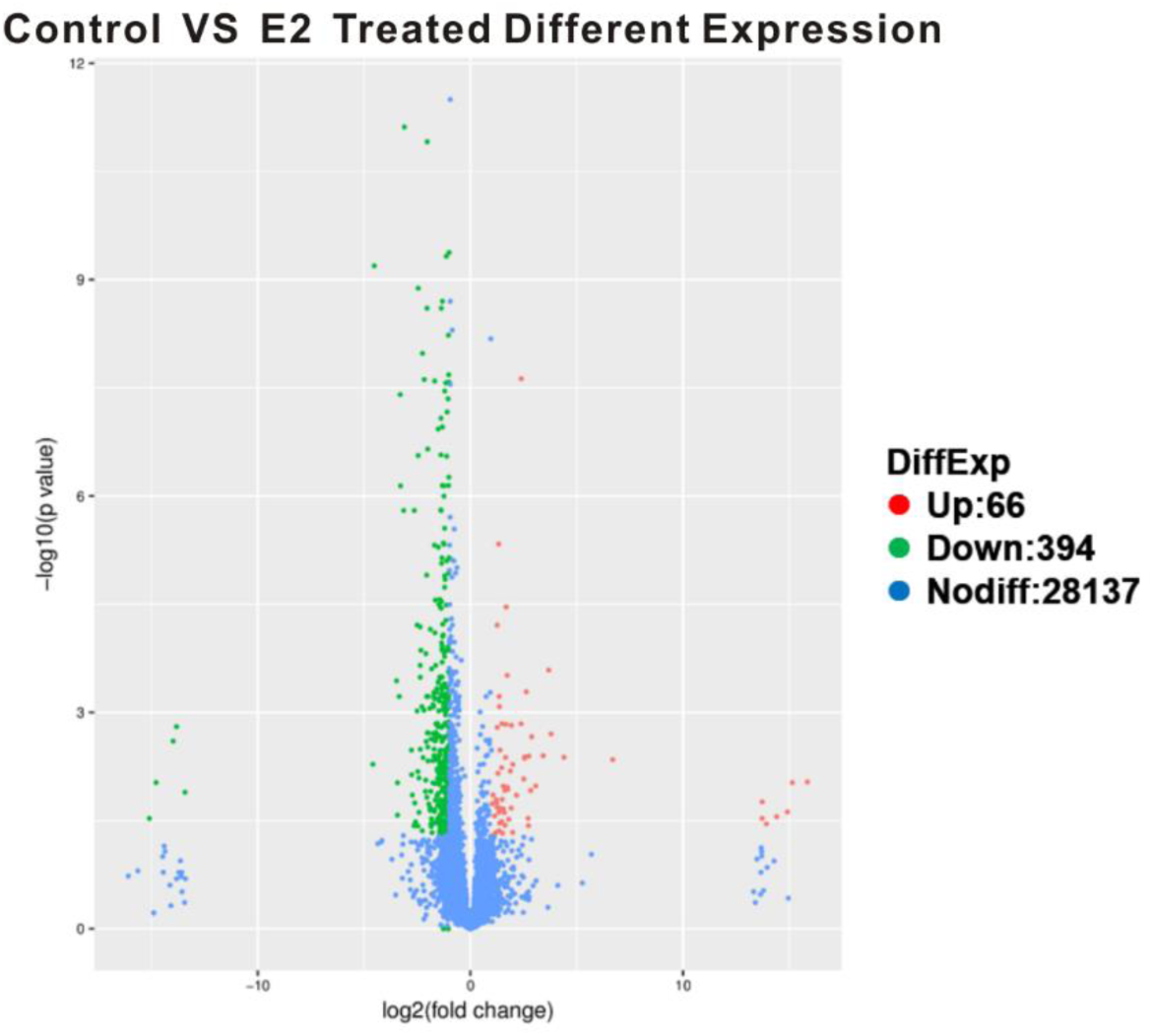
The RNA-seq identified DEGs. The screening criteria to identify significant differences in the expression of genes are |log2Ratio|≥1 and q-value≤0.05.

### GO analysis and KEGG pathway functional analysis of enriched DEGs

GO is an internationally standardized gene functional classification system that provides a set of dynamically updated vocabularies to comprehensively describe the properties of genes and gene products in organisms. The most enriched GO results for the three ontologies are shown in Table 2. We found that genes that affect biological regulation, metabolic processes, development, anatomical structure development, responses to stimulation, systems development, cell differentiation, cell communication, regulation of gene expression, and signal transduction were enriched in the DEGs (Table 2).

**Table 2.** Gene ontology enriched by differentially expression genes.

| Gene ontology enriched by differentially expressed genes (DEGs) |  |  |  |
| --- | --- | --- | --- |
| GO Id | GO-term | Hits | P-value |
| GO:0065007 | Biological regulation | 216 | 0.00455 |
| GO:0008152 | Metabolic process | 180 | 2.71E-06 |
| GO:0032502 | Developmental process | 118 | 0.00265 |
| GO:0048856 | Anatomical structure development | 103 | 0.00012 |
| GO:0050896 | Response to stimulus | 109 | 0.00265 |
| GO:0048731 | System development | 83 | 0.00326 |
| GO:0030154 | Cell differentiation | 73 | 9.09E-11 |
| GO:0007154 | Cell communication | 73 | 1.71E-06 |
| GO:0010468 | Regulation of gene expression | 98 | 0.00104 |
| GO:0007165 | Signal transduction | 62 | 9.34E-07 |

Multiple genes coordinate their biological functions in vivo. Pathway-based analysis enables further understanding of the biological functions of the genes. KEGG is the main public pathway database. Because we focus on the bone metabolism process, so bone metabolisms pathway was come into our view. We found that the DEGs were enriched for genes involved in the pathways for cancer, MAPK signaling, TGF-β signaling, focal adhesion, Wnt signaling, NF-κB signaling, and osteoclast signaling (Table 3). The TGF-β and Wnt signaling pathways are important in osteoblasts. Through the pathway maps, we identified alterations in the expression of *Bmpr1a*, *Tgfbr1*, *Rock1*, and *sp1*, which are involved in the TGF-β signaling pathway (Table 3). We also noted alterations in the expression of *Lrp5*, *Vangl2*, and *Rock1*, which are involved in Wnt signaling (Table 3). In the cell cycle signaling pathway, we noted alteration in the expression *Ccna2 Stag2*, *Atr*, *Smc1a* and *Rbl1.* These data show that the effects of estrogen maybe mediated via the TGF-β and Wnt signaling pathways in MC3T3-E1 cells.

**Table 3.** KEGG pathway enriched by differentially expression genes.

| Pathway ID | Pathway Terms | Hits Genes | P-value |
| --- | --- | --- | --- |
| ko05200 | Pathways in cancer | *Kras, Cul2, Appl1, Chuk, Hif1a, Msh2, Mecom, Birc2, Ccna2, Fgf5, Fos* | 0.001764262 |
| ko04350 | TGF-beta signaling pathway | *Sp1, Inhba, Tgfbr1a, Rbl1, Rock1, Sp1, Bmp1ra* | 0.004798816 |
| ko04010 | MAPK signaling pathway | *Kras, Chuk, Rasa1, Tgfbr1a, Mecom, Rasa2, Fgf5, Fos* | 0.01030574 |
| ko04510 | Focal adhesion | *Pdpk1, Vav3, Arhgap5, Birc2, Rock1, Actn2* | 0.01144867 |
| ko04310 | Wnt signaling pathway | *Lrp6, Vangl2, Rock1* | 0.01263819 |
| ko04064 | NF-kappa B signaling pathway | *Chuk, Birc2* | 0.01721087 |
| ko04380 | Osteoclast differentiation | *Chuk, Fos, Tgfbr1a* | 0.0313562 |
| ko04810 | Cell cycle | *Atr, Smc1a, Rbl1, Ccna2, Stag1,* | 0.04867427 |

### Confirmation of DEGs expression by qRT-PCR

We chose the candidate target genes from the bone metabolic signaling pathway. The candidate genes and their relative gene functions are listed in Fig. 3A. Among those genes, *Sp1*, *Tgfbr1*, and *Rock1* belong to the TGF-β signaling pathway; *Bmpr1a* belongs to the BMP signaling pathway; *Fos* is from the osteoclast differentiation pathway; *Lrp6 is* from the Wnt signaling pathway; and *Adra1b* belongs to the calcium signaling pathway. The qRT-PCR results indicated that *Bmpr1a* (Fig. 3B), *Tgfbr1* (Fig. 3C), *Rock1* (Fig. 3D), *Lrp6*, (Fig. 3F), and *Sp1*(Fig. 3H) were down-regulated by E2 treatment, whereas *Fos* (Fig. 3E) and *Adra1b* (Fig. 3G) were up-regulated. These results confirmed the differential expression after E2 treatment of the genes identified by RNA-seq.

**Figure 3.**
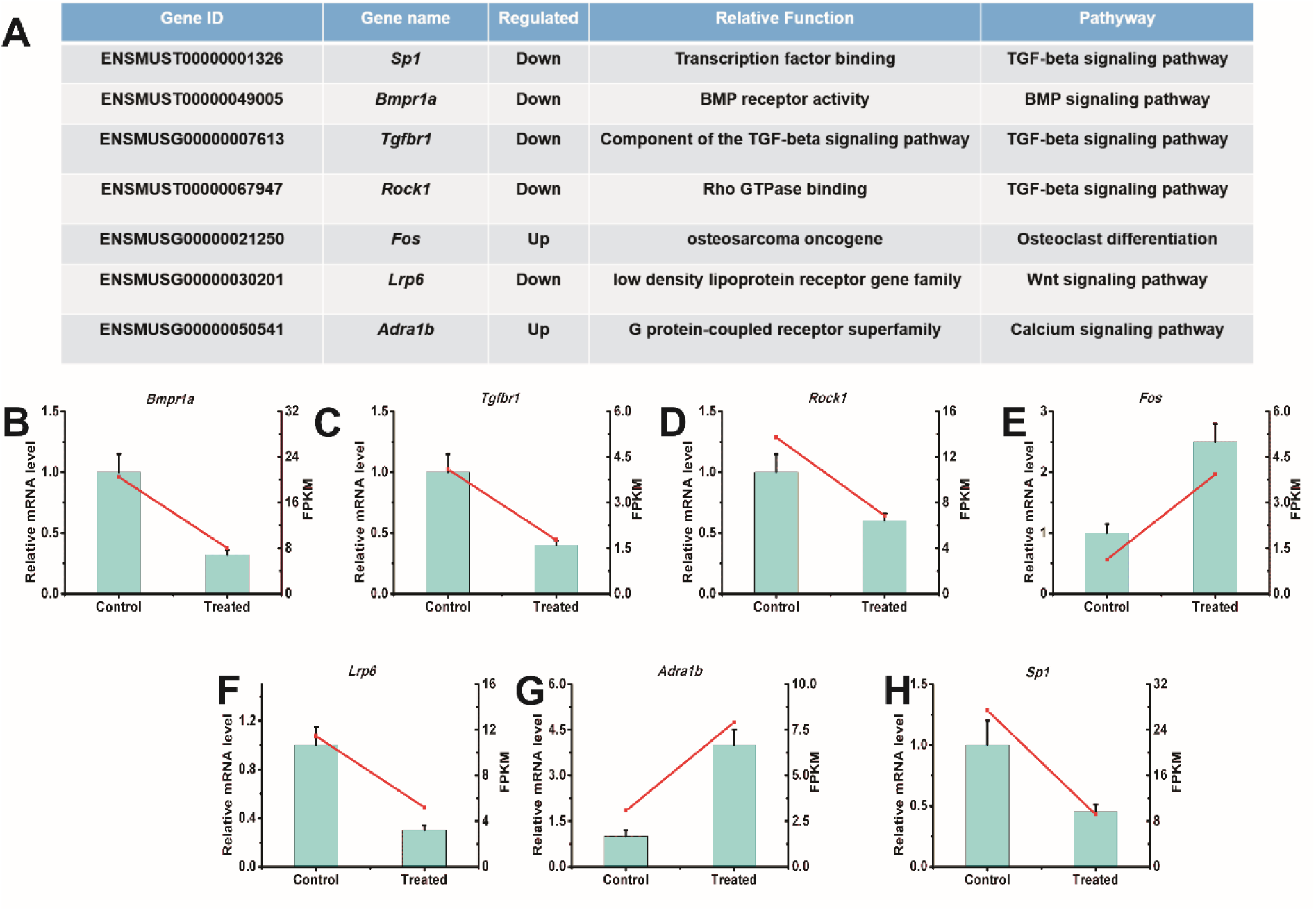
Confirmation of the mRNA levels of the DEGs by qRT-PCR. We treated MC3T3-E1 cells with 10 nM E2, then extracted total RNA for qRT-PCR. (A) The candidate genes involved in bone metabolism. (B) The expression level of *Bmpr1a.* (C) The expression level of *Tgfbr1.* (D) The expression level of *Rock1.* (E) The expression level of *Fos.* (F) The expression level of *Lrp6.* (G) The expression level of *Adra1b.* (H) The expression level of *Sp1.* Three independent experiments were conducted. Student’s t-test was performed. ^∗^, P < 0.05; ^∗∗^, P < 0.01.

### Estrogen negatively regulated gene expression via ERβ

Estrogen is an important endocrine regulator of bone metabolism, which mediates it effects via ERs [18]. ERs are transcription factors that bind EREs to activate or repress target gene expression [19]. Two mammalian ERs have been identified, ERα and ERβ [20]. Previous studies revealed that estrogen represses *SOST* expression via ERβ [21]. To investigate if the differential gene expression in the E2-treated MC3T3-E1cells was mediated via ERβ, we analyzed the *Bmpr1a* and*Tgfbr1* promoters using online bioinformatics software (http://jaspar.binf.ku.dk/). The results identified three EREs and two EREs in the *Bmpr1a* and *Tgfbr1*gene promoters, respectively (Fig. 4A). To determine if E2 repressed the candidate genes via a particular ER, we treated MC3T3-E1cells with 10nM E2, then added the ERα antagonist MPP or the ERβ antagonist PTHPP. We then performed qRT-PCR to investigate candidate gene expression. We found that E2 treatment significantly down-regulated *Bmpr1a* and *Tgfbr1*, compared to their expression in the DMSO control groups, in the MC3T3-E1 cells (Fig.4B-D). MPP did not rescue the E2-mediated down-regulation of *Bmpr1a* and *Tgfbr1* (Fig. 4B-D). However, PTHPP treatment rescued the down-regulation of *Bmpr1a* and *Tgfbr1* (Fig. 4B-D). Rescue of the candidate target gene expression by PTHPP, but not MPP, revealed that estrogen negatively regulates gene expression via ERβ.

**Figure 4.**
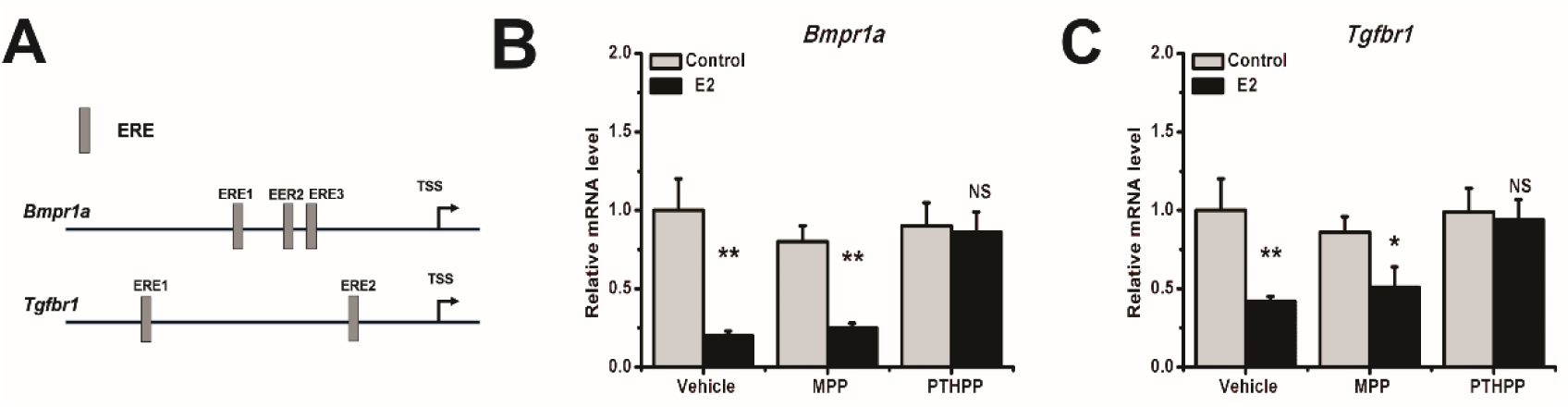
Estrogen negatively regulated gene expression via ERβ. (A) Promoter analysis for*Bmpr1a*, *Tgfbr1*, and *Lrp6.* T. MC3T3-E1cells were treated with 10nM E2, and 0.1μM MPP or 0.1μM PTHPP, then candidate target gene mRNA levels were quantified via qRT-PCR. (B) The expression level of *Bmpr1a* under different treatment conditions. (C) The expression level of *Tgfbr1* under different treatment conditions. Three independent experiments were conducted. Student’s t test was performed. ^∗^, P < 0.05; ^∗∗^, P < 0.01; NS, not significant.

### E2 increased ERβ binding to the EREs of the *Bmpr1a* and *Tgfb1* promoters in MC3T3-E1cells

We next addressed the potential functionality of the EREs located in the *Bmpr1a* and *Tgfbr1* promoters. We performed ChIP experiments using an ERβ antibody to determine if ERβ bound to the EREs in the *Bmpr1a* and *Tgfbr1* promoters. We found that ERβ bound to ERE1 and ERE3, but not ERE2, in the *Bmpr1a* promoter (Fig. 5A). Both of the EREs located in the *Tgfbr1* promoter were bound by ERβ (Fig. 5B). To investigate if E2 treatment affected the binding of ERβ in the *Bmpr1a* and *Tgfbr1* promoters, we treated the MC3T3-E1 cells with E2 for 72h, then performed ChIP, followed by qRT-PCR. E2 treatment increased ERβ binding to ERE1 and ERE3 in the *Bmpr1a* promoter (Fig. 6C), and ERE1 and ERE2 in the *Tgfbr1* promoter (Fig. 5D). These data demonstrated that ERβ can bind to the *Bmpr1a* and *Tgfbr1* promoters, and that E2 increased ERβ binding to the EREs of the *Bmpr1a* and *Tgfbr1* promoters in MC3T3-E1cells.

**Figure 5.**
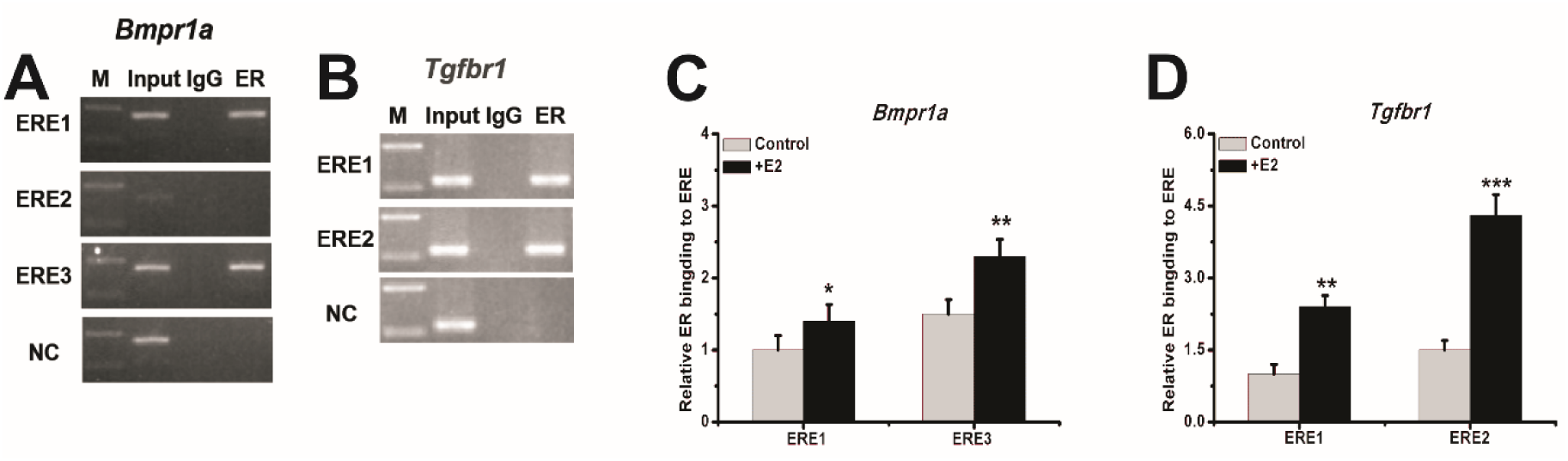
E2 increased ERβ binding to the EREs located in the *Bmpr1a* and *Tgfbr1* promoters in MC3T3-E1cells. We performed ChIP experiments on MC3T3-E1cells using an ERβ antibody. (A) ERβ binding to the ERE1 and ERE3, but not ERE2, located in the*Bmpr1a* promoter. (B) ERβ binding to the ERE1 and ERE2 located in the *Tgfbr1* promoter. We performed qRT-PCR after ChIP on cells treated with E2. (C) E2 increased ERβ binding to the *Bmpr1a* promoter. (D) E2 increased ERβ binding to the *Tgfbr1* promoter. Three independent experiments were conducted. Student’s t-test was performed. ^∗^, P<0.05; ^∗∗^, P<0.01.

## Discussion

E2, a synthetic, exogenous estrogen that is functionally similar to endogenous estrogen, is commonly used to study the function of estrogen. MC3T3-E1 cells are a newborn mouse calvarias cell line, which is commonly used in the study of osteoblasts. ERα and ERβ have also been identified in this cell line, so MC3T3-E1 can be used to study the role of estrogen in the regulation of osteoblast function.

We found that estrogen increases the rates of proliferation and apoptosis in MC3T3-E1 cells. We used CCK-8 and TUNEL assays to investigate the proliferation and apoptosis of MC3T3-E1 cells after E2 treatment. We found no significant differences in the groups treated with low concentrations of E2 after three days. However, high concentrations of E2 promoted the proliferation and apoptosis of MC3T3-E1 cells. E2-mediated increases in the proliferation of MC3T3-E1 cells have been reported in previous studies[22, 23], but we also found a slight but statistically significant increase in the number of apoptotic cells in our high concentration group, which indicated that treatment with E2 for three days was slightly toxic to MC3T3-E1 cells.

Estrogen affects many signaling pathways that are associated with bone metabolism. In both transcriptome libraries, these pathways can be identified by GO enrichment and KEGG pathway analysis of the DEGs. After E2 treatment of the MC3T3-E1 cells, cell differentiation-and cell cycle-related genes were significantly enriched among the DEGs. The involvement of these pathways can explain why E2-treated MC3T3-E1 cells undergo increased cell proliferation and apoptosis. KEGG signaling pathway analysis indicated that genes in the Wnt, MAPK, and calcium signaling pathways, and osteoclast differentiation genes, were significantly enriched among the DEGs. The enrichment of genes in these bone metabolic pathways among the DEGs demonstrated osteoblast-like differentiation of the MC3T3-E1 cells.

However, in our sequencing results, in addition to these signaling pathways, we found that the TGF-β signaling pathway and cancer-related signaling pathways were also enriched, in contrast with the research of Zhen et al. The TGF-β signaling pathway plays an important role in bone differentiation, especially in osteogenesis [24]. The TGF-β superfamily of ligands includes BMPs, growth and differentiation factors, anti-müllerian hormone, activin, nodal, and the TGF-β proteins [25]. Signaling begins with the binding of a TGF-β superfamily ligand to a TGF-β type II receptor [26]. The type II receptor is a serine/threonine receptor kinase, which catalyzes the phosphorylation of the type I receptor [25]. Each class of ligand binds to a specific type II receptor. In mammals, there are seven known type I receptors and five type II receptors [25]. In our sequencing results, we found that *Tgfbr1* and *Bmpr1a* levels were significantly reduced in MC3T3-E1 cells after E2 treatment, and qRT-PCR confirmed this result. *Tgfbr1*is also called activin receptor-like kinase receptor (*Alk5)* [27], ALK5 protein is strongly expressed in perichondrial progenitor cells for osteoblasts, and in a thin chondrocyte layer located adjacent to the perichondrium in the peripheral cartilage [28]. Conditional knockout of the TGF-β type I receptor *Alk5* in skeletal progenitor cells resulted in growth plates that had an abnormally thin perichondrial cell layer, as well as reduced proliferation and differentiation of osteoblasts [29].

The BMP receptors are a family of transmembrane serine/threonine kinases that include the type I receptors *Bmpr1a* and *Bmpr1b*, and the type II receptor Bmpr2 [25, 30]. Deletion of the receptor *Bmpr1a* in osteoblast lineage cells with a *Dmp1*-Cre caused a dramatic increase in trabecular bone mass in postnatal mice, which was due to a marked increase in osteoblast numbers. In addition to increasing the osteoblast numbers in the trabecular bone. [31]. Another study revealed that conditional deletion of *Bmpr1a* in differentiated osteoclasts negatively regulates osteoclast differentiation. However, osteoblast-specific deletion of *Bmpr1a* resulted in increased bone volume with marked decreases in BFR in tibias at eight weeks of age [32, 33].Thus, physiological *Bmpr1a* signaling in bone exerts a dual function in both restricting pre-osteoblast proliferation and promoting osteoblast activity. If the impact of decreased *Tgfbr1* and *Bmpr1a* expression on MC3T3-E1 cell differentiation or increasing osteoclast activity must be further explored.

Cancer-related signaling pathways are the second most abundantly enriched type of signaling pathway in our DEGs, and many types of cancers relative signaling are enriched, including colorectal cancer, pancreatic cancer, non-small cell lung cancer, endometrial cancer, and prostate cancer. This is consistent with clinical findings: estrogen is commonly used as a compensatory drug for estrogen loss, after long-term estrogen treatment estrogen in the initiation of many type cancer [34–36]. Although estrogen is an effective clinical treatment for postmenopausal osteoporosis and other estrogen deficiency-related diseases, its negative role cannot be ignored.

Estrogen inhibits the expression of *Tgfbr1* and *Bmpr1a* through ERβ. Previous studies have also shown that estrogen can inhibit the expression of *SOST* through ERβ [21]. To confirm that estrogen inhibits the expression of *Tgfbr1* and *Bmpr1a* through ERβ, we used ERα- and ERβ-specific inhibitors, and found that only the ERβ inhibitor rescued the expression of *Tgfbr1* and *Bmpr1a.* In addition, our ChIP experiment also demonstrated that ERβ binds to the *Tgfbr1* and *Bmpr1a* promoters. In mouse lung cancer tissue, estrogen was found to inhibit the expression of *Bmpr2* through ERα [37]. Estrogen also inhibits the activity of the TGF-β signaling pathway by promoting Smad2/3 degradation [38]. In another study, estrogen replacement decreases the accumulation of TGF-β1 on vascular smooth muscle cell (VSMC) [39]. In this study, we found that the ER signaling pathway and the TGF-β signaling pathway participate in crosstalk, and ERβ can inhibit *Tgfbr1.* Therefore, estrogen can down-regulate the expression of *Tgfbr1* and *Bmpr1a* in MC3T3-E1 cells.

## Conclusions

In conclusion, estrogen can affect bone metabolism via several bone metabolism-related signaling pathways. In MC3T3-E1 cells, estrogen appears to affect bone by negatively regulating *Tgfbr1* and *Bmpr1a* expression. Our research provides a new understanding of the mechanism by which estrogen acts on osteoblasts.

